# A lack of open data standards for large infrastructure project hampers social-ecological research in the Brazilian Amazon

**DOI:** 10.1101/2024.02.06.579211

**Authors:** J.L. Hyde, A.C. Swanson, S.A. Bohlman, S. Athayde, E.M. Bruna, D.R. Valle

**Affiliations:** School of Forest, Fisheries, and Geomatics Sciences, University of Florida, 136 Newins-Ziegler Hall, Gainesville, FL, 32611 USA.; Department of Global and Sociocultural Studies and Kimberly Green Latin American and Caribbean Center, Florida International University. Modesto A. Maidique Campus, Deuxieme Maison (DM) 353, 11200 SW 8th Street, Miami, FL 33199.; Center for Latin American Studies, University of Florida, 319 Grinter Hall, P.O. Box 115530, Gainesville, FL, 32611-5530, USA.; Department of Wildlife Ecology & Conservation, University of Florida, 110 Newins-Ziegler Hall, Gainesville, FL, 32611 USA

**Keywords:** open data, infrastructure, social-ecological research, conservation, tropics, Brazil, Amazon

## Abstract

New infrastructure projects are planned or under construction in several countries, including in the bioculturally diverse Amazon, Mekong, and Congo regions. While infrastructure development can improve human health and living standards, it may also lead to environmental degradation and social change. Accessible, high quality data about infrastructure projects is essential for both monitoring them and studying their social and environmental impacts. We investigated the availability and quality of data on infrastructure projects in the Brazilian Amazon by reviewing the academic literature and surveying researchers from the conservation and development community, then used the results of these surveys to identify critical data attributes for the gathering, organizing, and sharing of infrastructure data by social-ecological researchers and practitioners.

Although data on infrastructure in the Brazilian Amazon were generally available, they were often of poor quality and lacked information critical for monitoring and research. Data were often difficult to find and reformat, resulting in loss of time and resources for researchers and other stakeholders. Discrepancies between researchers’ survey responses on data needs and the types of data used in peer-reviewed articles on infrastructure projects indicate the following information was often missing: geographic extent of the project, construction and operation dates, and project type (e.g., paved vs unpaved road). Including these data in a standardized format, along with making them more readily accessible by hosting them in public repositories and ensuring they are current and comprehensive, would facilitate research and improve planning, decision-making, and monitoring of existing and future infrastructure projects in Brazil and other developing countries.

**Highlights:**

- Infrastructure projects across the globe spur economic development but also lead to social- ecological degradation.
- Researchers and practitioners need current and comprehensive data to better mitigate social- ecological changes from infrastructure.
- Infrastructure data are often unavailable, inaccessible, or incomplete.
- Finding and organizing datasets cost researchers hundreds of hours and may lead to abandoned projects.
- To promote better research outcomes, governments and NGOs should ensure datasets on infrastructure are accessible, current, comprehensive, and include such vital information as the project’s geographic extent, dates of construction and operation, project type, and essential technical data.

## 1. Introduction

Access to comprehensive, high-quality infrastructure project data is critical to studying, monitoring, and mitigating the social-ecological impacts of infrastructure (Joppa et al., 2016). This is particularly important give the millions of roads, dams, hydroways, ports, transmission lines and other major infrastructure projects that are currently operational, under construction, or planned worldwide, including planned massive regional, national, or multi-national infrastructure expansions (e.g., the Initiative for the Integration of the Regional Infrastructure of South America (IIRSA)^1^, and China’s Belt and Road Initiative^2^). Governments, nongovernmental organizations, and project funders collect and make available such data to monitor compliance and assess environmental impacts (Ciborra, 2005). Transparency and accountability resulting from making data available throughout the development process may help minimize inefficiency, corruption, and the mismanagement of public construction projects that have resulted in an annual loss of $4 trillion globally (Transparancy and Accountability Initiative, 2014). Researchers could help improve estimates of trade-offs and impacts for various project alternatives (Laurance et al., 2015), and third parties may bring innovative ideas and solutions to the table (Janssen et al., 2012). They can also hold the government accountable for including social-environmental variables in licensing or construction decisions and developing adequate consultation and compensations processes for affected populations (Moran et al., 2018; Pereira, 2021; Transparancy and Accountability Initiative, 2014). While the expansion and improvement of infrastructure can help increase standards of living and improve human health (Brenneman & Kerf, n.d.; Calderón & Servén, 2004; Estache, 2003; Johansson, 2002; Martínez & Ebenhack, 2008; Slough et al., 2015), large infrastructure projects can also lead to environmental degradation (Laurance, 2018; Laurance & Arrea, 2017; Pfaff et al., 2018; C. M. Souza et al., 2019), and negatively impact Indigenous peoples and other local communities (Arrifano et al., 2018; Fearnside, 1999; Gauthier & Moran, 2018). These social-ecological impacts cannot be properly identified and quantified without information about the infrastructure projects themselves.

Providing access to data and information about public or public-private infrastructure projects is typically the responsibility of a government institution, which usually also holds the responsibility for monitoring existing and planned projects for licensing or other administrative purposes and, thus, should have relevant information about these projects. Many countries have access to information laws requiring the release of information to the public (Kaufmann & Bellver, 2005; Relly, 2010) or have signed transparency pledges. Full implementation of these policies is rare, however, due in part to resource and technological constraints, lack of motivation and capacity among agencies, or unclear designation of responsibility (Attard et al., 2015; Ciborra, 2005; Di Ciommo, 2015; Janssen et al., 2012; Wang & Lo, 2016). Consequently, data about public or public-private infrastructure projects are often unavailable (Attard et al., 2015).

Even when infrastructure data are available, they may not be of a sufficient quality for specific social-ecological research. Task-independent data quality standards, which have been proposed by several entities, apply to datasets independent of research question or usage and focus on the completeness, accuracy, and currency of information (Open Data Charter, 2015; Pipino et al., 2002; Vetrò et al., 2016). These standards do not provide guidance on what information should be included within a dataset. However, for nearly all research projects, data quality is defined as the degree of usefulness in a particular task or context and is highly dependent on the user (Stvilia et al., 2007), equiring context-specific content. Consequently, even when data do meet task-independent quality standards, the dataset may still be of limited use because it lacks correct or sufficient information to guide a specific decision or to enable a specific task. For example, basic information about project location or date of construction is not always readily available to researchers, requiring them to invest significant time and resources searcher for or collecting these data (Hyde et al., 2018; Klarenberg et al., 2019; Tucker Lima et al., 2016).

Brazil is a culturally and ecologically hyperdiverse country, but major infrastructure development plans throughout the country, and especially in the Amazon, threaten this diversity (Athayde et al., 2019). Brazil has a strong legal framework to promote transparency, codifying the right to access information in the 1988 Brazilian Constitution and reinforcing this right with various national and international laws, ordinances, and supporting institutions^3^. In 2011, Brazil co-founded the Open Government Partnership (OGP)^4^, which seeks to promote transparency, empower citizens, fight corruption, and harness new technologies to strengthen governance. Unfortunately, most Brazilian government portals are not in compliance with international open government data criteria (Di Ciommo, 2015). It is also unknown whether data standards are followed throughout all sectors or if the standards are adequate for conducting meaningful research into social-ecological impacts of development.

Guidelines for the content of infrastructure datasets may improve the usefulness of these datasets for social-ecological research by ensuring they contain certain critical attributes and information (Joppa et al., 2016). Using infrastructure development in the Brazilian Amazon as a case-study, we conducted a systematic review of how infrastructure data have been used in social-ecological research in academic publications. Specifically, we asked: 1) What data are required for social-ecological research related to infrastructure projects? 2) How accessible and complete are public datasets on infrastructure projects in the Brazilian Amazon? We then surveyed practitioners and researchers about their data needs and efforts in searching for and using data on infrastructure. We used the results of our literature review and survey to identify what attributes should be included in all infrastructure datasets to maximize the utility of these datasets for researchers and other interested parties. Finally, we evaluated two datasets available on Brazilian government websites to determine how well they conformed to task-independent standards and whether they included the critical attributes we identified. While our study focuses on open data from Brazil, our recommendations are broadly applicable to infrastructure and development projects across the world.

## 2. Materials & Methods

### 2.1 Systematic literature review

We performed a systematic literature review to determine what information researchers have used when assessing the social and ecological impacts of infrastructure projects in the Brazilian Amazon. We only considered studies published after the passage of the Brazilian Federal Access to Information Law in 2011 (Lei. 12.527/201^5^). In accordance with this law, after 2011 scientists should have been able to acquire government data on infrastructure projects from Brazilian government agencies if this policy was fully enacted. We performed the literature review using the Web of Science’s^6^ “Core Collection” in September and October 2018. The search strings we used for the literature review are available at https://doi.org/10.5281/zenodo.10626908. We specifically looked for studies focusing on environmental impacts, management, or conservation in relation to current or planned infrastructure projects in the Brazilian Amazon. We only included studies if they specifically used some type of infrastructure dataset or information in their analysis or required infrastructure data to plan the research study. For each study, we determined the type of information used about infrastructure projects and focused on the project attributes (e.g., construction date, location, budget, etc.). We recoded the citation, topic, academic discipline, infrastructure type, the dataset(s) and types of data used, and the infrastructure attributes for each study.

### 2.2 Key informant survey

To understand data needs and experiences, we surveyed key researchers and practitioners who focus on social and/or ecological topics from a list of the corresponding authors of the papers in the literature review, members of the Amazon Dams International Research Network^7^ (ADN; Athayde et al., 2019), and members of the Governance and Infrastructure in the Amazon^8^ (GIA) working group (Mere-Roncal et al., 2021). The ADN and GIA coordinate social-ecological research and information-sharing about infrastructure in the Amazon and are comprised of researchers, NGO practitioners, and members of government agencies. WIth the survey, we collected demographic information and asked participants questions about types of infrastructure projects for which they searched, the information required about these projects, where they searched for information, how long it took to find relevant information and format it for use, what they did if they could not find appropriate data, and about data quality based on task-independent standards (Vetrò et al., 2016). Finally, we asked participants to list and rank infrastructure project attributes that were important for their use. This survey was approved by the University of Florida’s Institutional Review Boards (IRB #B201600928). The full survey is available at https://doi.org/10.5281/zenodo.10626908.

From the survey responses, we summarized which attributes were most important across infrastructure projects. We compared the data survey respondents wanted to data used in the literature and considered discrepancies between the two sources a possible data gap where necessary data might not be available. We evaluated the data quality and the amount of effort spent on data gathering, cleaning, and formatting by performing summary statistics on survey responses. To examine differences in data quality between data retrieved from government versus non-government sources, we only considered answers from participants who reported retrieving data exclusively from a government repository or exclusively from a non-government repository. We also combined responses for all non-government sources (i.e., academic, NGO, other).

### 2.3 Proposing critical attributes for infrastructure data sets

Based on attributes used in the literature review and survey participants’ rankings of attribute importance for informing social-ecological research and decision-making, we created context-specific standards for infrastructure datasets that are complimentary to the task-independent data quality standards. We considered attributes that were ranked in the top five more than 40% of the time as critical for inclusion in infrastructure datasets. By identifying these critical attributes, we strived to ensure the availability of information required to conduct social-ecological research about infrastructure projects.

### 2.4 Evaluating available infrastructure datasets

To further understand the quality of open data on infrastructure from the Amazon region, we evaluated two infrastructure datasets on whether they contained the attributes we identified as critical for social-ecological research and whether they complied with the task-independent framework provided by (Vetrò et al., 2016). Our test cases were large dams and roads as they are drivers of social-ecological change in the Amazon(Chen et al., 2015; Latrubesse et al., 2017; Laurance & Arrea, 2017; Nepstad et al., 2001) and frequently appeared in our survey responses and literature review. Therefore, it is especially important that these data are of high quality and useable for social-ecological research.

We downloaded data on May 22, 2019 from the agencies that oversee the dams and roads, the Agencia Nacional de Energia Elétrica (ANEEL)^9^ and the Departamento Nacional de Infraestrutura de Transportes (DNIT)^10^, respectively. We evaluated the quality of these two publicly-available infrastructure datasets based inclusion of critical information we identified and five characteristics from Vetrò et al.’s (2016) task-independent framework: (1) accuracy of spatial components, (2) completeness, (3) currency (up-to-date), (4) machine-readability, and (5) metadata quality. We assessed the spatial accuracy of the datasets by randomly selecting 50 existing projects in each dataset and verifying their locations in Google Earth using the same map projection. If the project was within 30 m (the size of a Landsat pixel) of the location listed on the dataset, it was considered spatially accurate.

We quantified how current the dataset was based on the date of the last update. Completeness was difficult to assess because it was unclear in many of the columns whether an empty cell was purposefully empty (the metadata did not provide this information). Instead of scoring the whole data set based on completeness, we chose the first two columns in each data set that were understandable without metadata and that clearly should have been complete, and we determined the percent of empty cells in these columns. Metadata quality was used as proxy for traceability (which measures the history of the data set) and understandability, both of which are somewhat subjective. Thus, we considered the metadata complete if it was present and contained explanations of the attributes in the data, its author, the geographic coordinate system of the shapefile, and the publication date.

## 3. Results

### 3.1 Systematic literature review

Sixty-two studies fit our criteria for the systematic literature review of articles that have investigated social-ecological impacts of infrastructure in the Brazilian Amazon. Together, the articles used infrastructure data 94 times (Figure 1A), requiring 236 attributes about those infrastructure projects (Figure 1B). Hydropower projects were the most common category of infrastructure investigated (43 datasets), followed by roads and highways (18 datasets). By far the most commonly used attribute about infrastructure projects was the geographic location of the project (66 times). The project name, its full geographic extent, and basic technical information were also used frequently. The articles focused on a range of topics, most frequently on social issues (such as displacement, livelihoods, socio-environmental conflict, human health, etc.), land use/land cover change, and aquatic ecology (Figure 1C).

**Figure 1.**
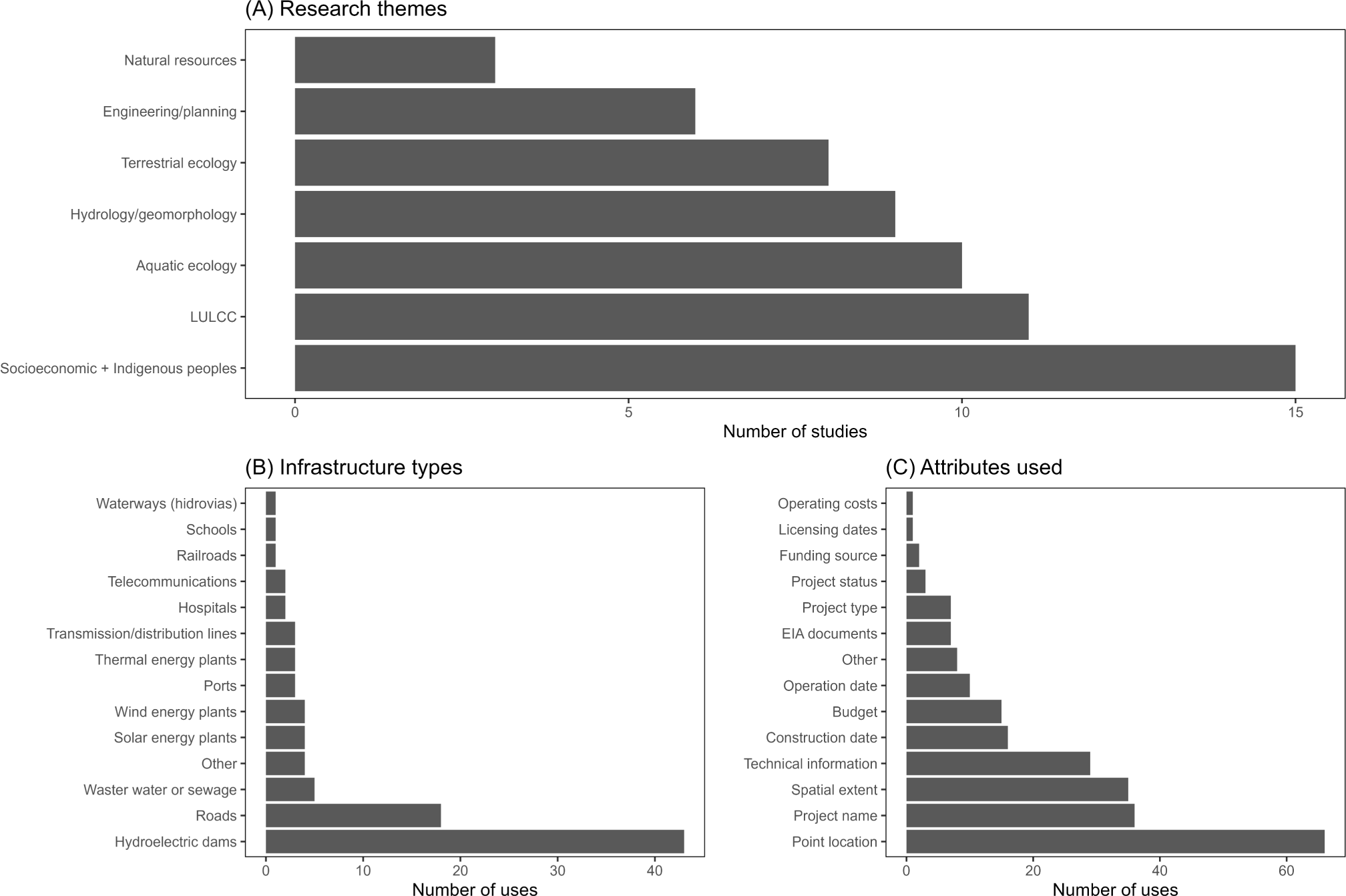
Number of articles in the systematic literature review conducted in the Web of Science database (WOS) for the 2011-2018 period grouped by (A) infrastructure project type researched; (B) data attributes used in the research (EIA stands for environmental impact assessment); and (C) thematic research area of the articles (LULC stands for land use/land cover)

### 3.2 Key informants survey

From the 472 people we contacted, a total of 87 people responded to the survey, with 68 completions (response rate = 18.4%, completion rate = 14.4%). Most participants (61.8%) were located in Brazil (Figure 2A) and 70.6% of respondents were in some stage of an academic career (Figure 2B). Participants were primarily researching socioeconomic topics (20.6%), land use/land cover change (20.0%), Indigenous peoples (16.3%), and natural resources (13.2%) (Figure 2C).

**Figure 2.**
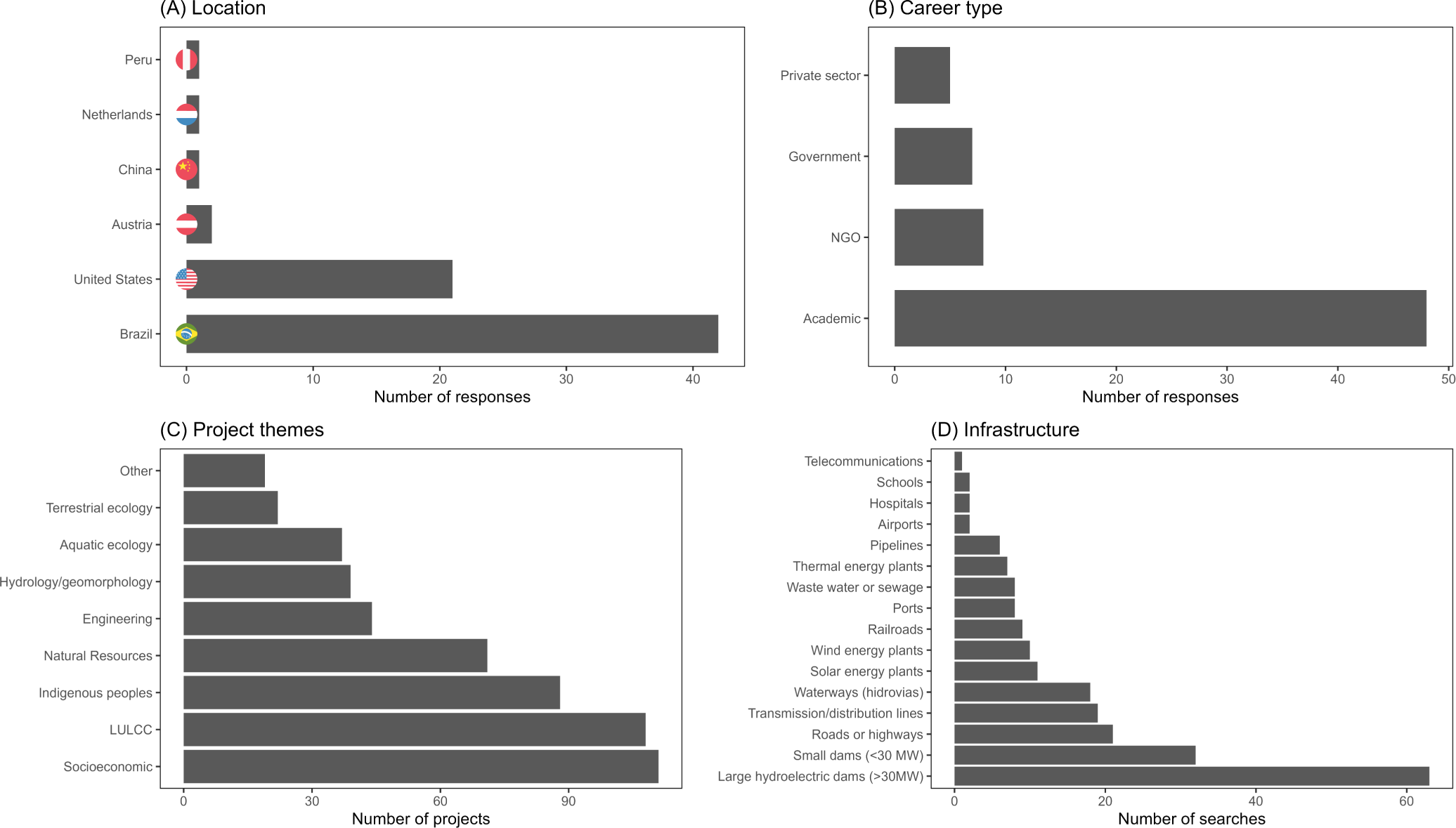
Survey responses for A) participants’ primary work locations and B) career type; C) themes of the projects for which participants needed infrastructure data. “LULCC” stands for land-use/land-cover change; D) types of infrastructure data searched for.

#### 3.2.1 High demand data sets

Collectively, the participants searched for infrastructure data for 539 projects. Combined, hydroelectric dams (28.8%), small dams (14.6%), and roads and highways (9.6%), accounted for more than half of the data searches (Figure 2D). Participants searched for information relatively evenly across project phases: 37.5% searched for information about the planning phase, 34.1% the construction phase, and 28.4% the post-construction phase. About half of searches (49.4%) were for datasets containing information about every project in the region from a specific infrastructure type (e.g. all large dams in the Amazon), 24.5% for information about two to five projects, and 26.1% for one specific project. Although participants searched for a wide range of information about the infrastructure projects; point location (10.5%), name (8.7%), status (8.4%), construction and operation dates (8.3% each), and full geographic extent (8.3%) were the most sought-after data attributes (Figure 3).

**Figure 3.**
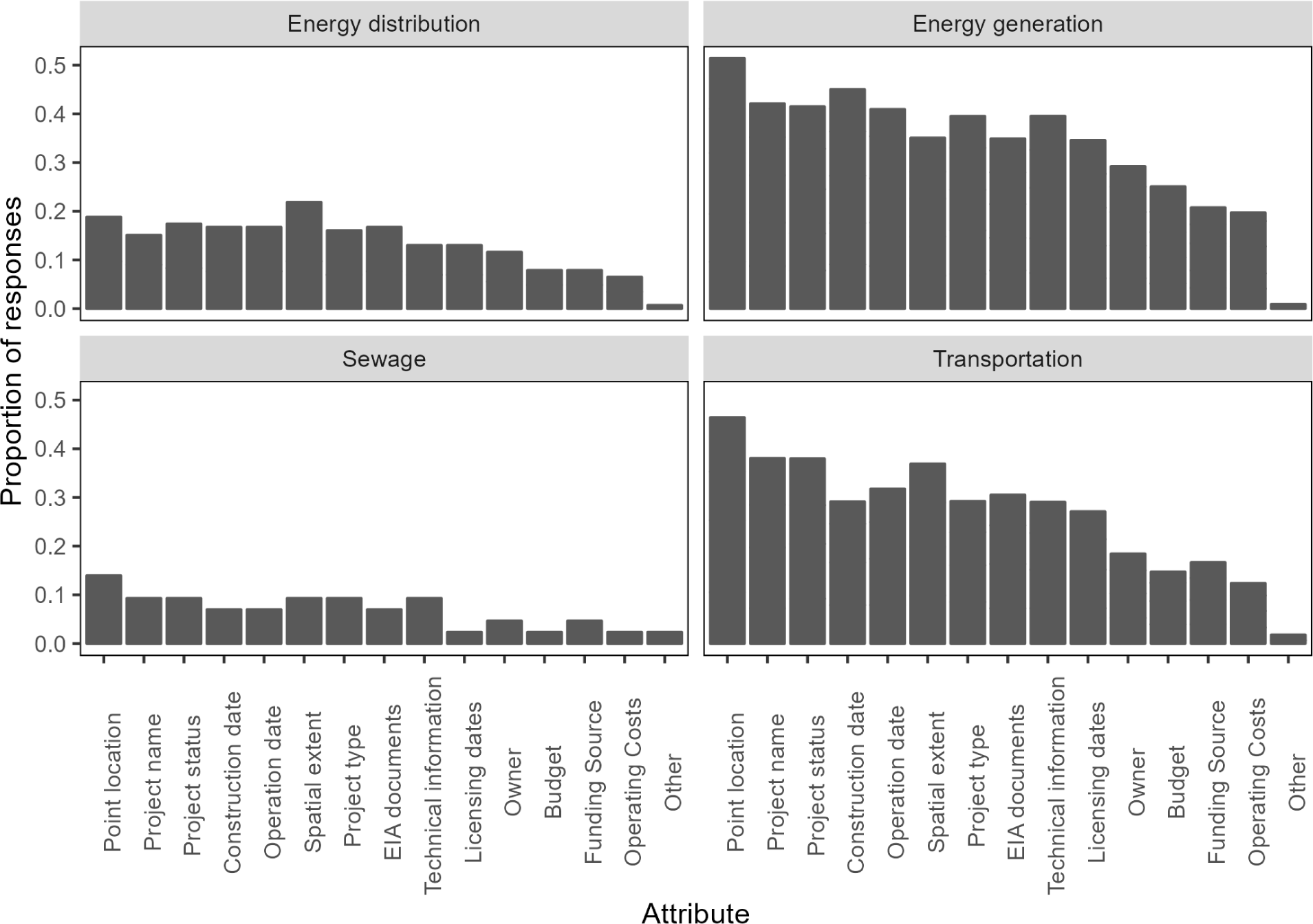
Proportion of searches by survey respondents for project attributes categorized by project type. Projects that had fewer than 10 attribute searches are not shown. Attributes are ordered according to their overall popularity from most to least frequently searched (left to right). Data were grouped into energy distribution (pipelines, transmission/distribution lines), energy generation (large and small dams, solar, thermal, wind), sewage (wastewater and sewage), and transportation (ports, railroads, roads and highways, and waterways/hidrovias).

There were substantial gaps between the attributes ranked within the top five by survey participants compared to the frequency of use these attributes in articles we reviewed (Figure 4). Most attributes were ranked withing the top five attributes required for social-ecological analysis in the survey at a higher rate than they were used in the studies we reviewed, including the geographic extent of the project, construction and operation dates, and project name and status. In contrast, point location was used more often in the literature than it was ranked in the top five attributes, possibly indicating this attribute was more available than the geographic extent of the infrastructure project, which may have been a more useful attribute. Combined, these results highlight potential gaps in infrastructure data availability.

**Figure 4.**
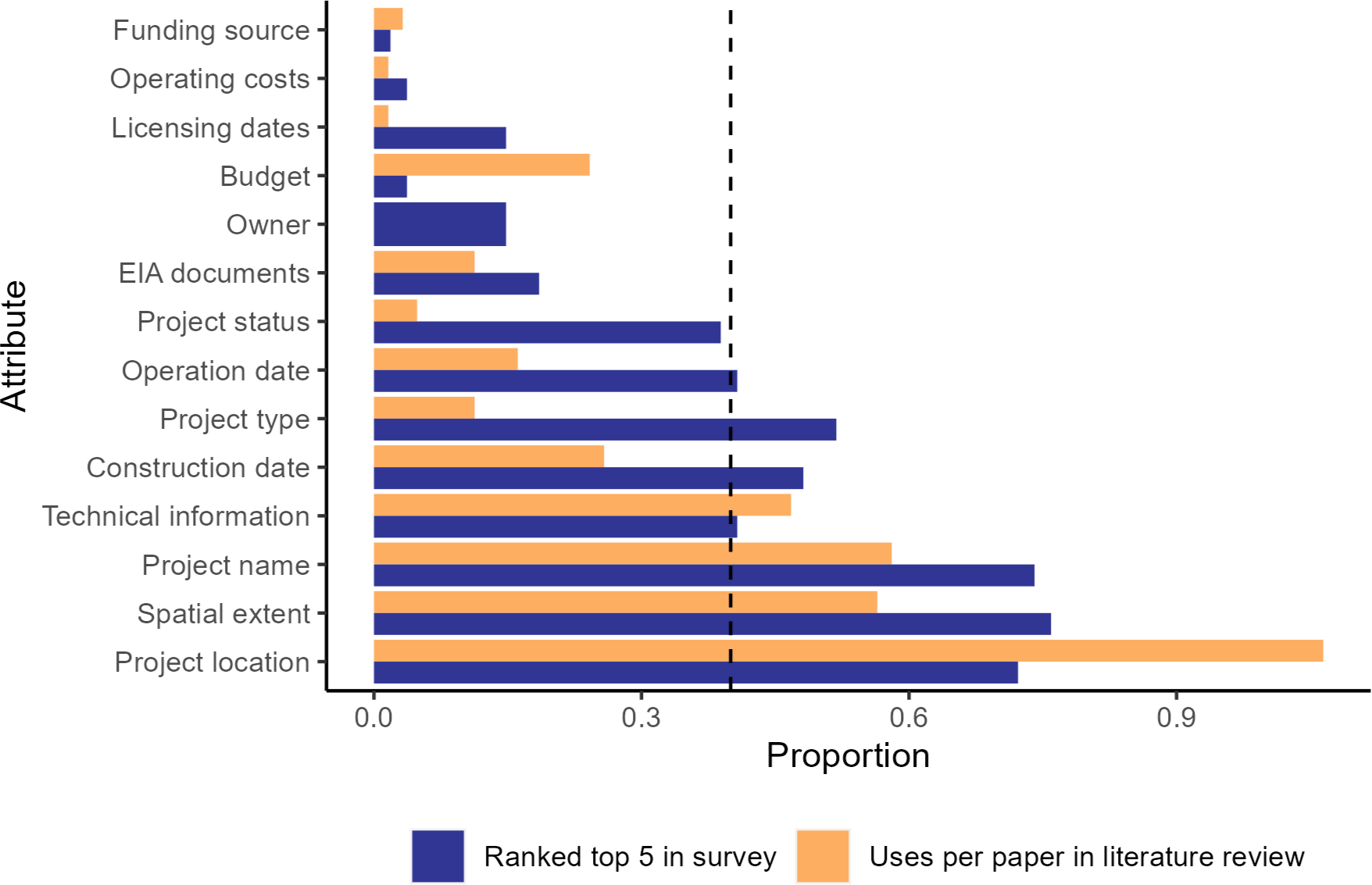
The proportion of times each attribute was used per paper in the literature review conducted in the Web of Science database (WOS) for the 2011-2018 period (orange), and the proportion of times each attribute was ranked in the top five (blue) for importance to include in an infrastructure data set by survey participants. EIA stands for environmental impact assessment. The dashed line at 0.4 is the cutoff for the attributes identified as critical for infrastructure datasets.

#### 3.2.2 Data accessibility and quality

Government sources were the most common place to search for information (39%), but academic and NGO sources were also frequently queried (31% and 24%, respectively) (Figure 5). Of the 188 searches for government data, 154 datasets were found from government sources, 27 from academic, NGO or other sources, and only seven were not found at all, highlighting the importance of universities, research institutions, and NGOs as providers of critical and reliable information on infrastructure projects. Respondents reported 147 searches for data from academic sources, of which 66% were found from these sources, 32% were found from alternate sources, and 2% were never found. There were 113 searches for data sourced by NGOs – 64% were found from NGO sources, 34% from other sources, and 2% were not found (Figure 5).

**Figure 5.**
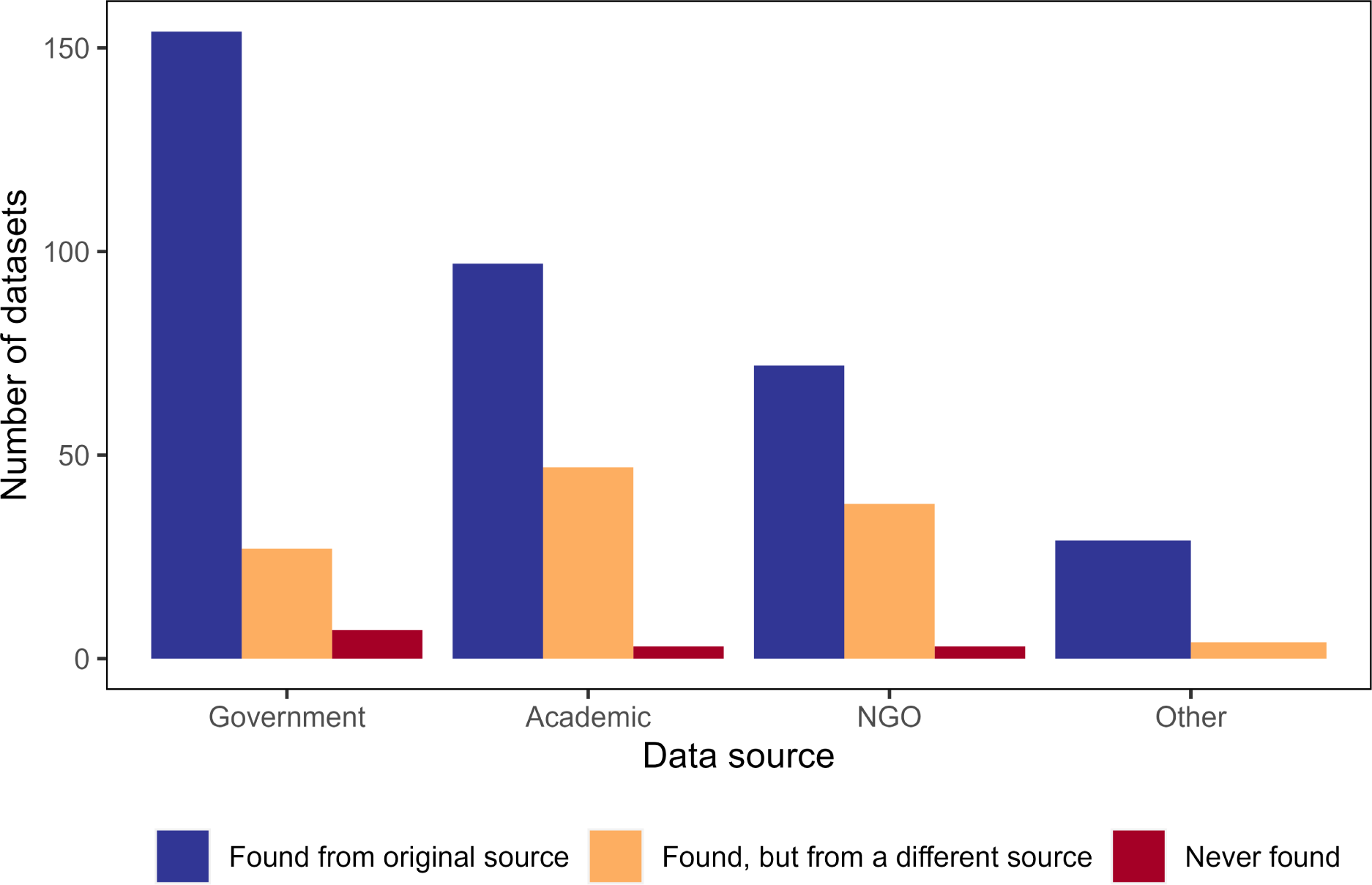
Number of infrastructure data sets survey participants searched for and found from the original search source (blue), searched for and found from a different search source (orange), or searched for but did not find (red).

From the surveyed participants’ responses, we assessed quality for datasets found from government and non-government sources (Figure 6). Participants reported high uncertainty about accuracy of non-spatial components of data from both government (65%) and non-government (63%) sources (Figure 6A). Government datasets were rated slightly lower in spatial accuracy than non-government datasets, but all participants reported that spatial accuracy was at least moderate in quality for both sources (Figure 6A). Data retrieved from government repositories was scored higher in task- independent standards that non-government data (Figure 6B). However, most datasets were still scored poorly for completeness (all cells filled), currency (updated within a year of the download date), and metadata quality (Figure 6B). Almost 75% of the government-sourced data was machine readable, compared to only 40% of non-government data (Figure 6B).

**Figure 6.**
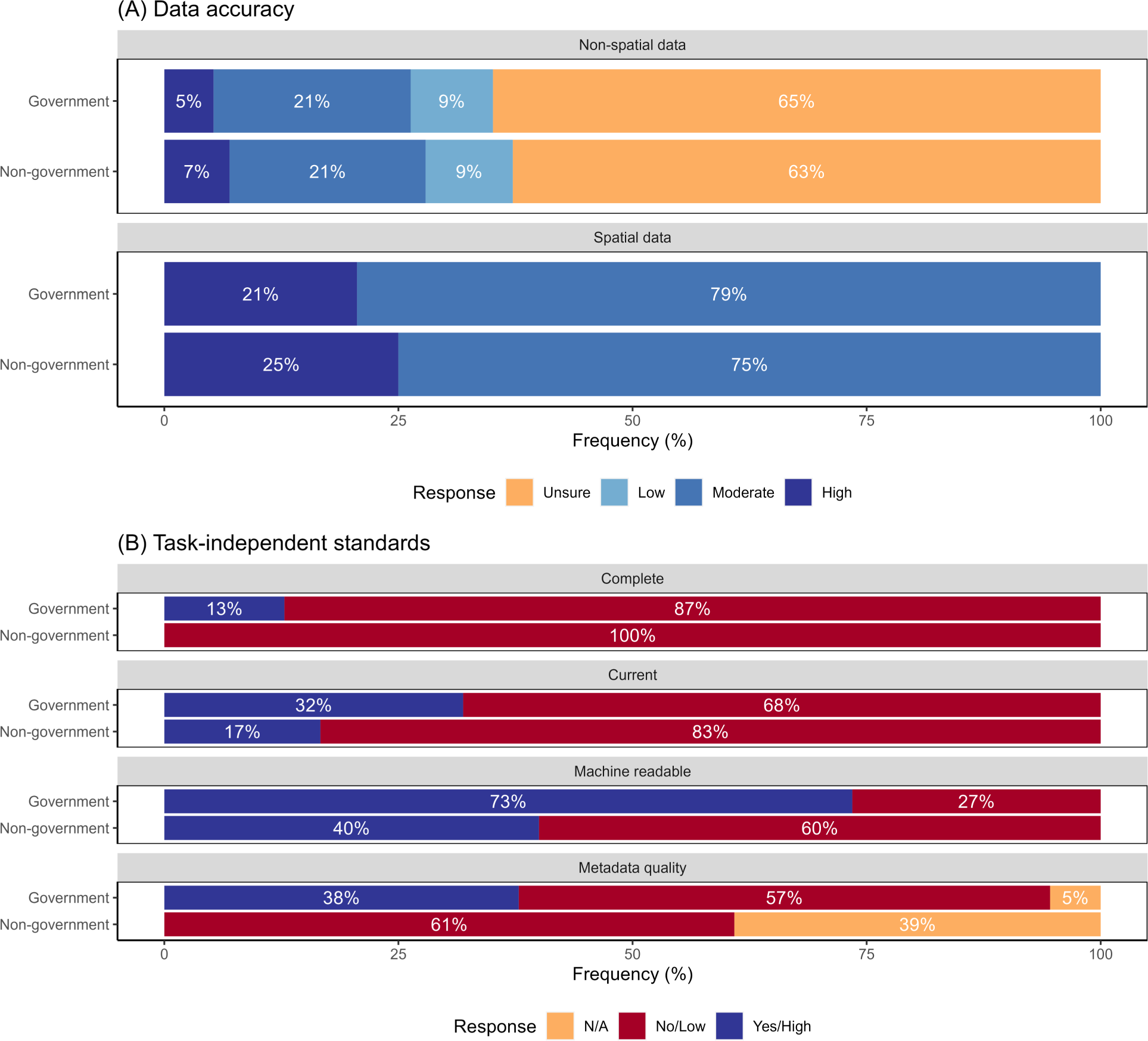
Responses to questions about data quality for datasets from government and non-government sources: (A) data accuracy (both spatial and non-spatial components); (B) task-independent standards – completeness, currency (< 1 year since update), machine readability, and metadata quality. N/A value for metadata quality indicates respondent did not require metadata for the dataset they used.

The majority of respondents (over 95%) were able to find data (Figure 7). Of those who found data, only 4% spent less than one hour searching for those data (Figure 7). One third of respondents reported spending between one and eight hours searching for data before finding it, while almost 30% spent more than 168 hours in their search for data (Figure 7). This is the equivalent of more than one month’s worth of work, assuming a 40-hour work week. Half of respondents who only found data from government sources spent more than eight hours searching for their datasets, while 44% of these respondents were able to find their data in one to eight hours (Figure 7). By comparison, 68% of respondents who found data only from non-government sources spent more than eight hours searching for data (Figure 7). However, this may be due to respondents who were unable to find data from government sources turning to non-government sources, requiring extra time to search for data. Ten respondents were unable to find the data they required (Figure 7) for a variety of infrastructure types: large hydroelectric dams (1), railroads (1), roads (1), solar energy plants (1), transmission/distribution lines (1), waterways (1), waste water/sewage (2), and small dams (2). Five of the respondents who were unable to find data abandoned their projects altogether, two used proxy datasets, two collected the data themselves, and a third respondent unsuccessfully attempted to collect data.

**Figure 7.**
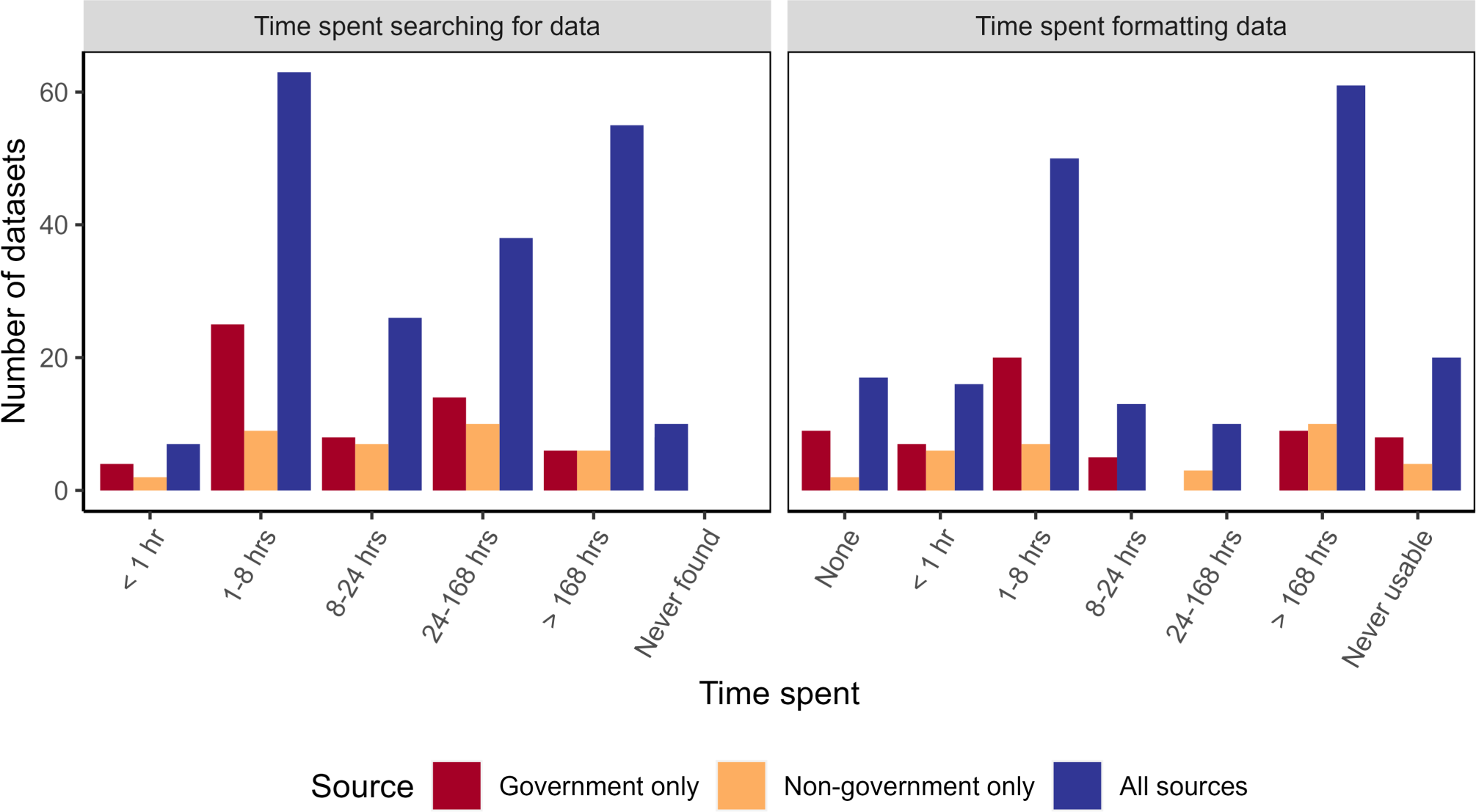
Time spent searching for and formatting data about large infrastructure projects in the Brazilian Amazon. Note that “all sources” represents the total number of datasets searched for or formatted and includes datasets found only from government or non-government sources and datasets found from both government and non-government sources.

Even when respondents were able to find data, 90% had to spend additional time putting those data into a usable format (Figure 7). More than half spent at least eight hours formatting data, and 37% spent at least 168 hours to make data useable (Figure 7). For those respondents who only used government data, 62% spent eight hours or less on data formatting, with more than 15% not spending any time on data formatting (Figure 7). However, another 15% respondents who only used government data had to spend more than 168 hours on data formatting to make use of their data (Figure 7). Only 47% of respondents who did not use government data spent eight hours or less on data formatting, while over 31% of these respondents spent more than 168 hours on this task (Figure 7). Twenty respondents reported never being able to get their data into a usable format, eight of whom were only using government data. As a result of data being unusable, seven respondents abandoned their projects. Of those respondents who continued with their projects, one used a proxy dataset and five collected new data. The unusable datasets varied in quality. Three were incomplete, eight were not current, four were not machine readable, and six had low quality metadata. Spatial accuracy across all unusable datasets was either high (3) or moderate (6), but 60% of respondents reported uncertainty about non-spatial data accuracy. No respondents found the non-spatial data for datasets that were unusable to have high accuracy.

### 3.3 Critical attributes for infrastructure datasets

Here, we propose critical attributes that should be included in all infrastructure datasets (Box 1). We based this proposal attributes that were ranked within the top five by at least 40% of respondents (Figure 4). While our survey explicitly described two geographical features of infrastructure datasets (point location and full geographic extent of the project boundaries), we included only the full geographic extent as a critical attribute because both attributes were ranked almost equally by survey respondents and because point location can be extracted from the full geographic extent. These attributes are the minimum information that should be readily accessible for all infrastructure projects to facilitate social-ecological research on impacts of these projects. In addition to including these content-specific attributes, data managers must also ensure that datasets are current, complete, and accurate, and provide these data in a machine-readable format with complete metadata.

**Box 1.**
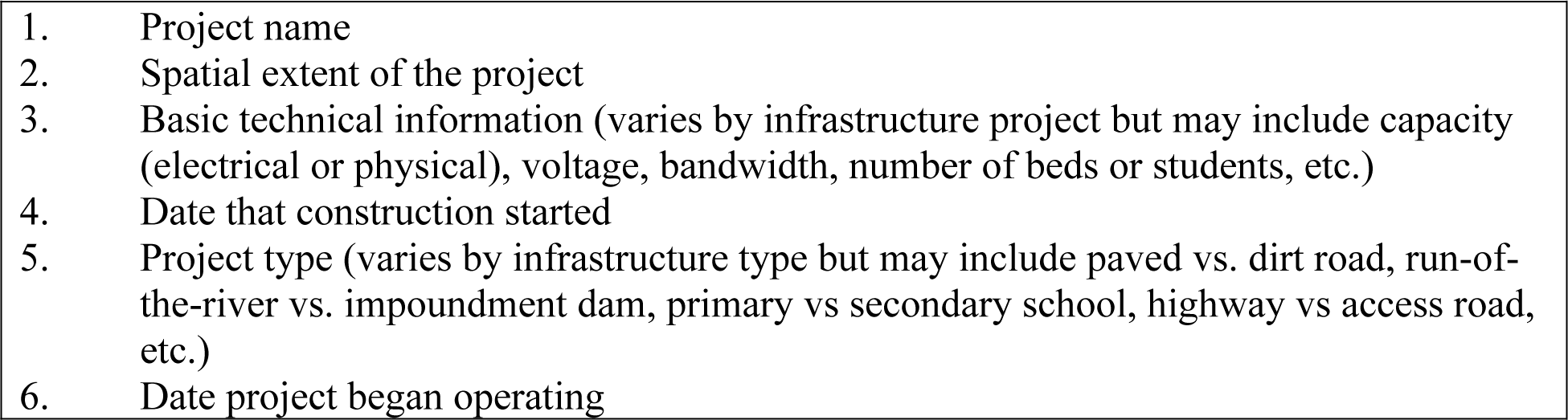
Critical attributes that should be included in all infrastructure datasets

### 3.4 Evaluation of public datasets

We evaluated whether two publicly available infrastructure datasets (large hydroelectric dams and roads) contained the critical attributes we proposed should be included in all infrastructure datasets and assessed the quality of these datasets based on the task-independent standards. Though we were not able to find either dataset on Brazil’s central data repository (dados.gov.br), the datasets were available from the websites of the agencies that oversee dams (Agencia Nacional de Energia Elétrica) and roads (Departamento Nacional de Infraestrutura de Transportes). Neither dataset contained all of the critical attributes we proposed (Table 1). While both datasets did include project names, neither included project construction or operation dates. The dams dataset included point data and technical information on reservoir sizes but did not include the geographic extent of reservoirs or the dam buildings. The roads dataset included project type (e.g., paved/unpaved, federal/state) and geographic extent in the form of line features for the full length of roads but lacked technical information.

**Table 1.** The quality and content of publicly available infrastructure data sets based on our proposed critical attributes and task-independent standards.

|  | Federal roads | Large dams |
| --- | --- | --- |
| Dataset origin | Departamento Nacional de Infraestructura de Transportes | Agencia Nacional de Energia Eléctrica |
| Critical attributes |  |  |
| Project Name | Yes | Yes |
| Geographic extent | Yes | No (point location only) |
| Basic technical info | No | Yes (capacity, drainage area) |
| Construction date | No | No |
| Project type | Yes (pavement status, federal status, road size) | No |
| Operation date | No | No |
| Task-independent standards |  |  |
| Spatially accuracy | 98% | 96% |
| Completeness (2 columns) | 100 % | 94.3% |
| Currentness | No information | Yes (updated 5/8/19) |
| Machine readable | Yes (shapefile, KMZ) | Yes (shapefile, KMZ) |
| Metadata |  |  |
| Author | No | No |
| Date of creation/update | No | Yes |
| Attribute descriptions | No | No |
| Datum | No | No |

The datasets had high spatial accuracy with 96% (dams) and 98% (roads) of randomly-selected points falling within 30 m of their location based on satellite imagery. To measure completeness, we determined the number of filled cells within the first two easily understandable columns: project name and owner for dams, project name and status for roads. For the dams dataset, 94.5% of cells within these two columns were filled while 100% of the cells within the two roads columns were filled. Across all columns, 17.5% (dams) and 21.7% (roads) of cells were missing data, but it was unclear whether these cells were supposed to be empty because there were no attribute descriptions in the metadata. The dams dataset was current, but there was no information on the currency of the roads dataset (Table 1). Both datasets were available in machine readable formats (ESRI shapefile or KMZ). The metadata quality was low for both datasets. The roads dataset included no metadata for author, date of creation or latest update, attribute description, or geographic datum. The metadata for the dams dataset only included information about the date of latest update (Table 1).

## 4. Discussion

### 4.1 Critical data attributes for social-ecological research

Public access to high quality data about current and planned infrastructure projects is critical for researching social-ecological impacts, conserving biodiversity, and human rights initiatives related to these projects. Our key informants survey and literature review revealed that the infrastructure project’s name, geographic extent, construction and operation dates, project type, and basic technical information are critical data for social-ecological research; these attributes, at minimum, should be included in openly-available infrastructure datasets improve their usability. This information should be made available for all current infrastructure. For planned infrastructure projects, which will not have exact construction and operation dates or spatial extents, datasets should include the proposed dates and area of the project. Furthermore, these data should conform to basic task-independent standards, namely being current, complete, accurate (spatially and non-spatially) and machine readable with informative metadata.

The data attributes we propose should be included in all publicly-available infrastructure datasets are used in many ways in social-ecological research. For example, Swanson and Bohlman (2021) used the names, operations dates, dam types, and geographic extent of reservoirs to quantify changes in land cover in the Tocantins River watershed after the installation of multiple hydropower dams. Nickerson et al. (2022) compared deforestation surrounding large hydropower dams and small dam clusters in the Legal Amazon, which required information about the construction and operation dates as well as technical information and type of dams. From an energy planning perspective, de Faria and Jaramillo (2017) investigated alternatives to hydropower expansion in the Amazon using location and capacity data on all current and planned wind, solar, thermal, and hydropower in the region.

Additionally, de Souza et al. (2023) used data for highways, waterways, railways, and ports, including project type, geographic extent, and technical information, to model port hinterlands in the Brazilian Amazon. Finally, Menezes et al. (2018) modeled vulnerability of Amazonian municipalities to climate change using technical information about hospitals. These studies demonstrate the importance of information about specific infrastructure projects to monitor social-ecological change related to new development and to improve planning and monitoring efforts for future infrastructure development.

They also illustrate that the content-specific attributes recommended for inclusion in infrastructure datasets are ubiquitous across infrastructure types.

Though this study focused on the Brazilian Amazon, the information needed from infrastructure data is likely to be consistent independent of location. For instance, de Faria and Jaramillo’s study (2017) went beyond the Brazilian Amazon and looked at the potential for renewable energy generation across Brazil. Likewise, Lupinetti-Cunha et al. (2022) and Tisler et al. (2022) both used the geographic extent and project type to model the effects of roadless areas on land cover change and conservation in Brazil. Beyond Brazil, Flecker et al. (2022) used location and technical information about hydropower dams to investigate how environmental impacts from damming could be reduced. Baird et al. (2021) investigated downstream impacts of dams and the need for Indigenous and traditional knowledge to mitigate those impacts in the Amazon, Canada, Laos, and Vietnam. They used information including dam names, operation and construction dates, spatial extent, and technical information to conduct their study. Finally, Ding et al. (2022) used information about project type to model the carbon emissions of 5G cell stations in China. These studies illustrate that the information identified in our literature review and through our key informants’ survey is applicable beyond the geographic boundaries of the Brazilian Legal Amazon. Though we developed these recommendations based on research conducted in the Brazilian Amazon, our recommendations were designed to be general enough to be applicable to infrastructure development initiatives across the globe and to improve the usability of infrastructure data for a broad range of research initiatives.

### 4.2 Data quality and availability for the Amazon

While we found that infrastructure data for projects in the Legal Amazon were generally available, finding them often required extensive, time-consuming searches. Furthermore, most were often low quality based on task-independent standards, which required users to spend additional time on data munging. The federal government is often the main regulator, if not the main funder and co-owner, of these large-scale infrastructural projects, and government repositories were the most common place where stakeholders searched for these data. Therefore, it is especially important that government agencies provide high quality data for projects under their jurisdiction. Unfortunately, our survey reveals that not all government data is easily accessible, thereby requiring that users spend significant time both searching for and formatting data. Studies of publicly available data from non-governmental sources have shown similar issues in accessibility and usability (Roche et al., 2015; Vines et al., 2014).

Although usefulness is context dependent, ensuring that data conforms to task-independent standards would likely alleviate some of this time input (de Oliveira & Silveira, 2018). Task-independent standards require data to be current, machine readable, complete, accurate, and accompanied by informative metadata. However, these standards do little to alleviate the issue of accessibility. Much of the time participants spent searching for data may have been due to difficult-to-navigate websites or because the Brazilian central repository^11^ does not host all the necessary data, so individuals may have had to search across government websites and jurisdictions (de Oliveira & Silveira, 2018). A central repository to host infrastructure data, either provided by the federal government or by non-government institutions, would likely reduce the search time thereby increasing the accessibility of information. For example, non-government central data repositories like Global Forest Watch or MapBiomas store some infrastructure data in addition to environmental data and could serve to collate data across jurisdictions.

These results demonstrate the costs of poor accessibility and quality. After previously investing time into searching for data (and in some cases, putting them into usable formats), poor quality or missing data resulted in 12 abandoned projects. Alternatively, researchers invested time and resources to collect new data, thereby having less time and resources for other projects. The cost of poor quality or inaccessible data in time and financial resources are more easily measured, but it is challenging to determine the costs associated with failing to generate potentially crucial information for assessing and managing impacts and planning future projects more sustainably. Delays or failures to create this information could have long-term consequences during the global infrastructure boom.

### 4.3 Gaps in data availability

There was a discrepancy between the attributes frequently requested by survey respondents and attributes that were frequently used in past scientific studies. Attributes that were ranked high in the survey but not used often in the literature review could reflect the accessibility gap for information that is not readily available. For example, construction and operation dates ranked high enough to be included in our list of critical infrastructure data attributes but were used in a lower proportion of studies. Similarly, point location was used far more often than the geographic boundaries of the project, although they were requested equally in the survey, likely indicating that the full geographic extent of projects is less available. This discrepancy highlights important data gaps that may hinder research.

Basic information such as the date of construction and the full spatial extent of the infrastructure are critical in tracking change related to infrastructure projects. The discrepancy between the desired attributes (from the survey) and previously used attributes (from the literature review) can indicate an important accessibility gap, with project type, construction and operation dates, and project status being the least accessible but highly demanded attributes.

### 4.4 Accuracy of assessed data

We reviewed the quality of public data sets for dams and roads from government websites. While Brazil appears to follow the laws regarding access to information by freely providing basic information about infrastructure projects, the overall task-independent and conservation-specific quality could be substantially improved to increase the usefulness of these data. Neither of the datasets we reviewed contained all the critical data we identified in our study, sometimes lacking data on a project’s full geographic extent, construction and operation dates, technical information and project type. Both had low-quality metadata that made it difficult to interpret many of the attributes. While both were spatially accurate, the dams data set did have the full geographic extent of either the dam buildings or the reservoirs. We were unable to come up with a reasonable way to tell if the dams and roads data sets we evaluated were complete or if the non-spatial components were accurate. For example, it is not clear that the datasets contained all the infrastructure projects in a given category (e.g. all the federal roads). It was also impossible to determine if the non-spatial data about the infrastructure projects are accurate without exhaustively searching planning documents, a task which would have taken dozens to hundreds of hours to complete. This uncertainty is also reflected in the frequent “I don’t know” responses from our survey participants about the accuracy of non-spatial components. It is easier, albeit time consuming, to validate the spatial accuracy of spatial data. However, judging the completeness of a spatial dataset requires field surveys or detailed reviews of satellite imagery or other remote sensing data. Hyde et al. (2018) hand-digitized all the transmission lines in the Legal Amazon and compared them to two public datasets. They found that the two datasets differed from each other, and only one was spatially accurate. Additionally, neither dataset included every transmission line in the region, though this was not immediately apparent without manually validating every satellite image for the Legal Amazon against the datasets, a process that took Hyde et al. (2018) almost six months.

This demonstrates the importance of external validation of spatial and non-spatial accuracy to ensure data quality. In fact, a system that allowed users to rate the completeness and accuracy of datasets could increase confidence in public datasets and reduce the time spent validating data before use.

### 4.5 Recommendations for improving data accessibility

In cases where governments have not or are unable to provide accessible and high-quality data, the research and NGO communities should strive to fill in these gaps. For example, data initiatives like RAISG, MapBiomas, Global Forest Watch, and SERVIR host infrastructure data. However, individual researchers should also be held responsible for sharing data that they collect on public archives. In fact, the academic community should encourage more journals to make the dissemination of data a requirement for publication, and grant funders should also require comprehensive data management and sharing plans for their grantees (Reichman et al., 2011). Models for data management and sharing plans already exist through the National Science Foundation (NSF) and the National Aeronautics and Space Administration (NASA). A culture of open data will reduce the redundant collection of data while allowing researchers to be credited for their work through citations. Furthermore, this will serve to improve data accessibility beyond only infrastructure data and reduce gaps in environmental and social data, which is equally important for monitoring social-ecological change.

### 4.6 Caveats

While considerable effort was made to obtain a representative sample of stakeholders interested in all infrastructure types in the Amazon region, the survey population was biased toward a hydropower focus. The hydropower bias may have resulted from sampling some of the survey respondents from the Amazon Dams International Research Network. However, we note that the literature review results also were also skewed towards dams. Thus, this may simply reflect the general focus of the scientific community on dams in the Amazon region in the context of the hydropower boom (Athayde et al., 2019). Despite the focus on dams, at least one person searched for every attribute for every infrastructure type, so we believe that our results and the critical data attributes can be used broadly across infrastructure types. Finally, most of our survey respondents held academic positions, so these results may reflect the needs of the academic research community more strongly than those of government, the private sector, or NGO communities.

### 4.8 Conclusions and future directions

Access to high quality data about infrastructure projects has the potential to improve the quality and efficiency of social-ecological research and assessing impacts related to new and planned development projects. With access to the best data available, third parties and governments can ensure the accuracy and accountability of environmental impact assessments (Laurance et al., 2015), fair compensation to impacted communities, and appropriate mitigation plans. They can also inform improved watershed-level planning, as opposed to the project-by-project planning (Athayde et al., 2019), as well as better identify projects that are more harmful than helpful. Importantly, this transparency can also potentially reduce corruption in the infrastructure sector (Kaufmann & Bellver, 2005; Ruijer et al., 2017). o achieve this, governments and third parties should release data sets that conform to task-independent standards and contain the critical attributes identified in this study. The research and government communities should also strive to remove barriers to accessibility by investing in comprehensive and up-to-date central repositories to host this data. As countries continue to expand and update their infrastructure, promoting transparency and data sharing about the projects is an important step in implementing the right to access to information, as well as improved public participation in decision-making related to current and future infrastructure.

## Acknowledgments

We would like to thank the University of Florida Water Institute, the School of Forest Resources and Conservation and the UF Tropical Conservation and Development Program (TCD) for their support, as well as Dr. Stephen Perz, Dr. David Kaplan, and the 2015 Water Institute Graduate Fellows. Dr. Charles Jekel was instrumental in helping with coding issues. We are grateful to our colleagues in in the Amazon Dams International Research Network/Rede Internacional de Pesquisa em Barragens Amazônicas/ Red Internacional de Investigación en Represas Amazónicas (ADN/RBA/ RIRA), to the participants of the civil society initiative and working group GT Infraestrutura, and to our associates at the Agência Nacional de Energia Elétrica (ANEEL) and Empresa de Pesquisa Energética (EPE) for their input and support during this research process. The survey was performed under the IRB #B201600928.

## Author Contributions

**J.L. Hyde:** Conceptualization, methodology, software, validation, formal analysis, investigation, resources, data curation, writing – original draft, writing – review & editing, project administration; **A.C. Swanson:** software, validation, formal analysis, data curation, writing – review & editing, visualization, project administration; **S. Bohlman:** resources, writing – review & editing, supervision, funding acquisition; **S. Athayde:** writing – review & editing, supervision, funding acquisition; **E.M. Bruna:** writing – review & editing, supervision; **D.R. Valle:** conceptualization, methodology, writing - review & editing, supervision, funding acquisition

## Data Availability Statement

Data created for this paper are available at https://doi.org/10.5281/zenodo.10626908. Code created for analyzing these data is available at https://zenodo.org/doi/10.5281/zenodo.10627391.

## Footnotes

1 http://www.iirsa.org/

2 http://english.gov.cn/beltAndRoad/

3 https://www.right2info.org/recent/access-to-public-information-in-brazil-what-will-change-with-law-no.-12.527-2011

4 www.opengovpartnership.org

5 Brasil. Lei ordinária n° 12.527, de 18 de novembro de 2011. Regula o acesso a informações previsto no inciso XXXIII do art. 5°, no inciso II do § 3° do art. 37 e no § 2° do art. 216 da Constituição Federal. Diário Oficial da União 2011; 18 nov.

6 The Web of Science (WOS), previously known as Web of Knowledge, is an online subscription-based scientific citation indexing service that provides a comprehensive citation search. The Web of Science Core Collection consists of six online databases: Science Citation Index; Social Sciences Citation Index. Arts and Humanities Citation Index; Emerging Sources Citation Index; Book Citation Index; and Conference Proceedings Citation Index. Additional databases available in WOS searches include SciELO Citation Index; BIOSIS Citation Index; MEDLINE1; CABI; and Zoological records. Website: https://clarivate.com/products/web-of-science/ Source: Wikipedia: https://en.wikipedia.org/wiki/Web_of_Science,

7 http://amazondamsnetwork.org

8 https://giamazon.org

9 https://sigel.aneel.gov.br/Down/

10 https://www.dnit.gov.br/planejamento-e-pesquisa/dnit-geo

11 http://dados.gov.br/

